# AsCas12a Exhibits Intrinsic, DNA-Independent ATPase Activity

**DOI:** 10.1101/2025.06.05.658205

**Authors:** Supreet Bhattacharya

## Abstract

Cas12a (previously Cpf1) is a class 2 CRISPR-Cas effector protein with RNA-guided DNA endonuclease activity and is widely used for genome editing. While its DNA cleavage and target recognition mechanisms have been studied extensively, the possibility of auxiliary enzymatic functions remains underexplored. Here, I report that *Acidaminococcus* sp. Cas12a (AsCas12a) possesses intrinsic ATPase activity, despite lacking canonical nucleotide-binding or hydrolysis motifs. Using a radiometric thin-layer chromatography (TLC) assay, I demonstrate that AsCas12a hydrolyzes ATP in a concentration and time-dependent manner. Importantly, this activity occurs independently of DNA cofactors, as neither single-stranded nor double-stranded DNA could influence the rate or extent of ATP hydrolysis. Bioinformatic analyses using NsitePred and SwissDock, as well as ATPbind, suggests potential ATP-binding residues with predicted favorable binding energies. This preliminary finding uncovers a previously unrecognized biochemical property of AsCas12a.

## 1. Introduction

Cas12a (or Cpf1) is a type V-A CRISPR-associated endo- and exonuclease that has garnered significant attention for its programmable DNA cleavage ability in genome editing and diagnostics [1–3]. Like Cas9, Cas12a is guided by a CRISPR RNA (crRNA) to introduce double-stranded breaks at target sites. While its structural domains and DNA targeting functions have been well characterized [4–6], auxiliary enzymatic activities associated with Cas12a remain relatively unexplored. In general, ATP hydrolysis plays critical roles in the function of many DNA-processing enzymes, including helicases, translocases, and remodeling factors [7–9]. However, Cas12a lacks conserved nucleotide-binding motifs such as Walker A or B, raising the question of whether it could still exhibit ATPase activity through non-canonical mechanisms. Previous studies on CRISPR-associated proteins have uncovered unexpected enzymatic functions, prompting us to investigate whether AsCas12a could similarly possess cryptic enzymatic activities. In this study, I used a radiolabeled ATP hydrolysis assay to test AsCas12a’s ability to catalyze ATP turnover in the presence and absence of DNA cofactors. The ATPase activity of the AsCas12a protein was discovered serendipitously-when in one of the assay to test for *E. coli*. RecA activity, it was kept as a control (10). Further, bioinformatics analysis suggested potential ATP-binding residues in AsCas12a, which needs to be validated further through other experiments. Overall, the findings reveal a DNA-independent ATPase activity intrinsic to AsCas12a, suggesting potential auxiliary roles beyond its nuclease function.

## 2 Materials and Methods

### 2.1 Purification of AsCas12a

AsCas12a was purified as previously described (ref.1 and Figure S1).

### 2.2 Preparation of DNA substrates

The sequences of oligonucleotides (ODNs) used in this study are listed in Table 1. ODNs were 5′ end-labelled with [γ-^32^P] ATP using T4 polynucleotide kinase, and DNA substrates were made as described previously (Table 2, ref. 11).

**Table 1.**
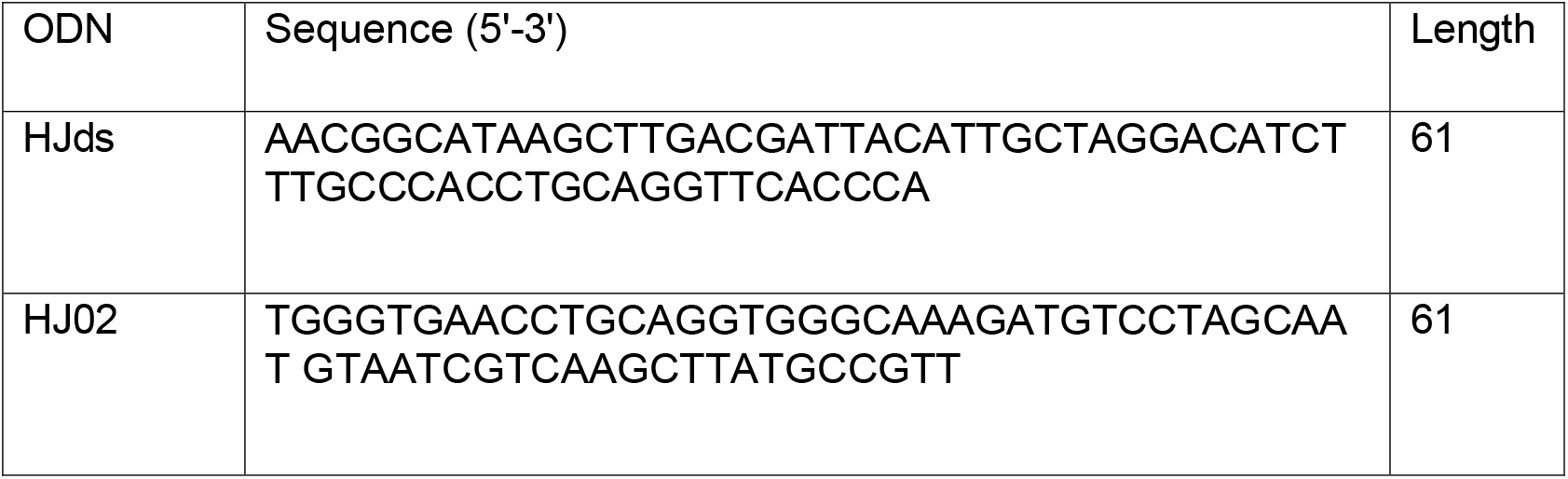
The sequences of oligonucleotides used to make various DNA substrates.

**Table 2.** Construction of synthetic DNA substrates by annealing respective oligos (see Methods in main text). Asterisks (*) indicate the ^32^P labeled strand.

| Substrate | Oligonucleotides used |
| --- | --- |
| Single-stranded DNA | HJ02* |
| Double-stranded DNA | HJ02* + HJds |

Oligonucleotide sequences for making DNA substrates-

### 2.3 ATPase assay

The assay was performed as previously described (11). Reaction mixes (20 µl), which contained 20 mM Tris-HCl buffer (pH 7.5), 12 mM MgCl_2_, 2 mM DTT, 15% glycerol, 100 µg/ml BSA, 200 µM cold ATP, 200 pM [γ-^32^P] ATP and indicated concentrations of AsCas12a, were incubated at 37 ºC for 1 h. The reaction was stopped by adding EDTA to a concentration of 25 mM. After incubation at 37 ºC for 10 min, aliquots (2 µl) of the reaction volume were spotted on a polyethyleneimine (PEI)-cellulose thin-layer plate (Whatman), and the reaction products were separated using a solution of 1 M formic acid and 0.5 M LiCl. The plate after air-drying, was exposed to a phosphorimager screen, and visualised using Fuji FLA-9000 laser scanner. The band intensities were analysed and quantified using UVI-BandMap (version 97.04). The data was plotted using Graph-Pad Prism software (version 5.0).

### 2.4 Bioinformatics analysis

The amino acid sequence of AsCas12a was retrieved from the Protein Data Bank using the PDB ID, 5B43. This sequence, formatted in FASTA, was then input into the NsitePred tool to identify potential nucleotide-binding residues. Further, the most probable binding site was subject to molecular docking using SwissDock. In a parallel analysis, the sequence file of AsCas12a formatted in FASTA was input into the ATPbind tool to identify potential ATP-binding residues/pockets.

## 3 Results

### 3.1 AsCas12a hydrolyzes ATP in a concentration and time-dependent manner

Despite lacking canonical ATP-binding motifs or domains typically required for ATP hydrolysis, AsCas12a unexpectedly exhibited ATPase activity when included as a control in assays for RecA (2, 10). To systematically test for intrinsic ATPase activity, purified wild-type AsCas12a was incubated with [γ-^32^P]ATP and monitored hydrolysis via TLC. Separation of substrate and product allowed for the visualization of free ^32^Pi release. AsCas12a catalyzed ATP hydrolysis in a concentration and time-dependent manner, with increasing amounts of ^32^Pi correlating with enzyme concentration up to 480 nM and time-points up to 50 min(Fig. 1-D). These data indicate that AsCas12a is capable of catalyzing ATP turnover efficiently.

**Figure 1.**
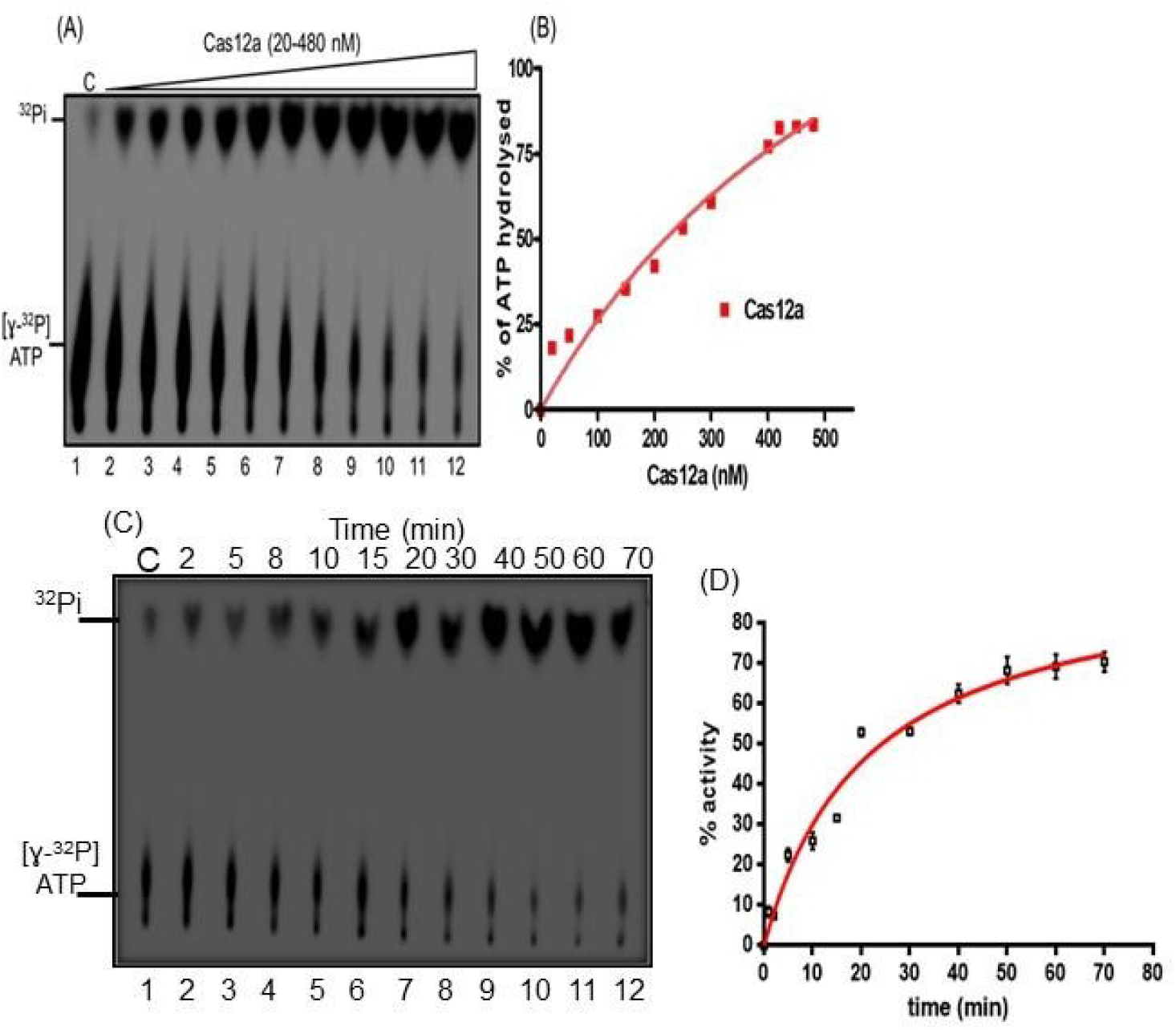
AsCas12a catalyzes hydrolysis of ATP to yield ADP and inorganic phosphate. (A), Representative image showing the separation of [γ-^32^P] ATP (substrate) and ^32^Pi (product) by thin-layer chromatography (TLC) after incubation with indicated concentration of AsCas12a (lanes 2-12). Control lane 1 lacked AsCas12a. The assay was performed as described in Material and methods section. Varying concentrations of AsCas12a is depicted by an empty triangle above the gel image. (B), Graph shows the amount (%) of ATP hydrolyzed versus increasing concentration of AsCas12a. (C), (D), Kinetics of ATP hydrolysis reaction mediated by AsCas12a and its quantification. The data shows mean ± S.D.from three independent experiments. Note that in the first lane of both panels (A) and (C), [γ-^32^P]ATP undergoes spontaneous degradation releasing Pi, and the products of ATP hydrolysis by AsCas12a migrate on TLC at same level.

### 3.2 ATPase activity of AsCas12a is independent of DNA cofactors

For many bacterial proteins, the property of ATPase activity is tightly linked to DNA metabolism, while in some others these are independent (12, 13). To determine whether DNA enhances or regulates ATP hydrolysis, the assay was repeated in the presence of either single-stranded DNA (ssDNA) or double-stranded DNA (dsDNA). The protocol involved pre-incubation of DNA with AsCas12a at 37 ºC for 10 minutes, followed by carrying out the ATPase assay. No significant difference was observed in the rate or total extent of ATP hydrolysis with either cofactor (single-stranded or double-stranded DNA) compared to the DNA-free condition (Fig. 2A–D). This indicates that AsCas12a’s ATPase activity is DNA-independent and likely intrinsic to the protein.

**Figure 2.**
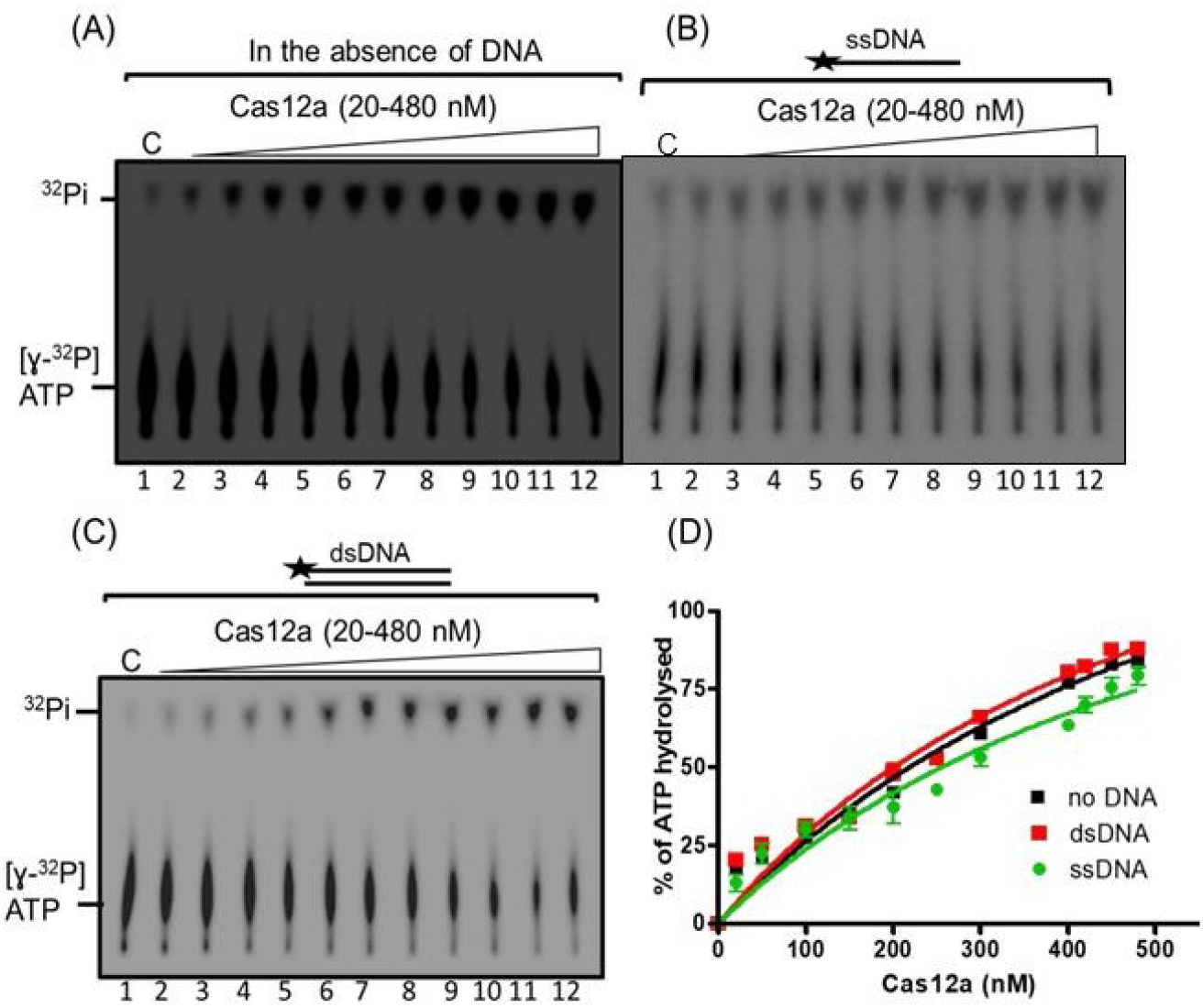
AsCas12a’s ATPase activity does not depend on DNA cofactors. TLC-based ATPase assays were performed in the absence (panel A, lanes 2-12) or presence of ssDNA (panel B, lanes 2-12) or dsDNA (panel C, lanes 2-12) and varying amounts of AsCas12a as described in Materials and methods section. (A-C), control lane 1 lacked AsCas12a. The empty triangles above the gel images indicate increasing concentrations of AsCas12a. (D) Graph shows quantitative comparison of ATPase activity (shown in panels A-C) in the presence of different DNA cofactors. The amount of ATP hydrolyzed was quantified, and the data were plotted as the percentage of ATP hydrolyzed versus increasing concentrations of AsCas12a. Each bar is the mean ± standard error of three independent experiments.

### 3.3 Predicting AsCas12a’s potential nucleotide-binding residues

Given the robust ATP hydrolysis activity exhibited by AsCas12a, it was pertinent to investigate its potential nucleotide-binding residues. The amino acid sequence of AsCas12a was obtained from the Protein Data Bank (PDB ID: 5B43) and analyzed using NsitePred, a freely accessible web-based tool that integrates sequence-derived structural features, evolutionary profiles, and alignment data to predict nucleotide-binding residues with high accuracy (14). NsitePred identified four ATP-binding residues, along with additional sites potentially involved in binding other nucleotides, including ADP, AMP, GTP, and GDP (Table 3). Notably, residues Pro1069, Ala1070, and Tyr1072 were predicted to constitute an ATP-binding site, which may function as a catalytic triad. These residues, all located within the NUC domain, were designated as the region of interest for molecular docking. Using SwissDock, ATP was docked as a ligand within this region (15) (Figure 3). The observed negative AC and SwissParam scores indicated favorable binding energy and strong predicted affinity (Table 4). In a parallel analysis, ATPbind identified two ATP-binding pockets, and notably, residues Arg696, Gln700 (within PAM-interacting domain), and Ser1068, His1071 (within NUC domain) were predicted with high confidence to constitute two ATP-binding sites (Figure 4 and Table 5) (16). Nonetheless, further validation through molecular dynamics simulations and site-directed mutagenesis will be required to confirm the relevance of the predicted ATP-binding site/s.

**Table 3.**
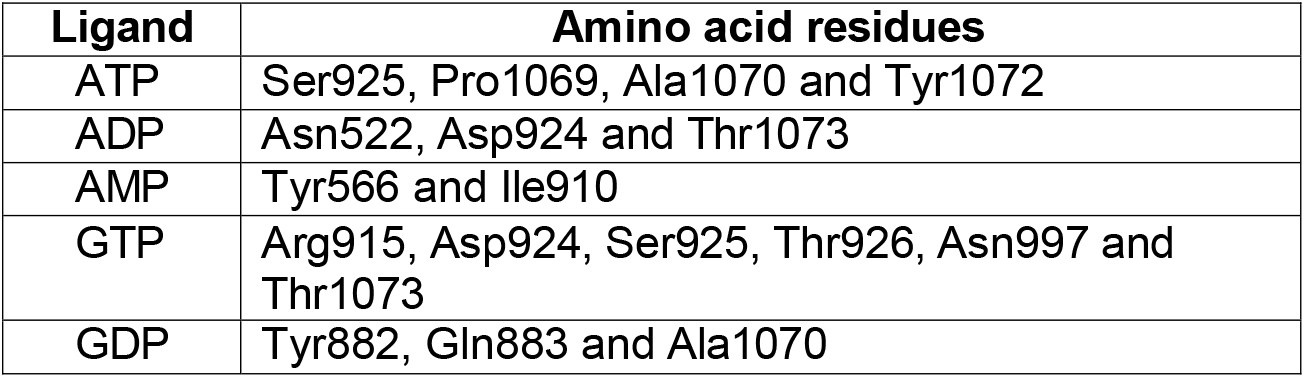
Possible nucleotide binding sites of AsCas12a according to NsitePred.

| **Ligand** | **Amino acid residues** |
| --- | --- |
| ATP | Ser925, Pro1069, Ala1070 and Tyr1072 |
| ADP | Asn522, Asp924 and Thr1073 |
| AMP | Tyr566 and Ile910 |
| GTP | Arg915, Asp924, Ser925, Thr926, Asn997 and Thr1073 |
| GDP | Tyr882, Gln883 and Ala1070 |

**Table 4.**
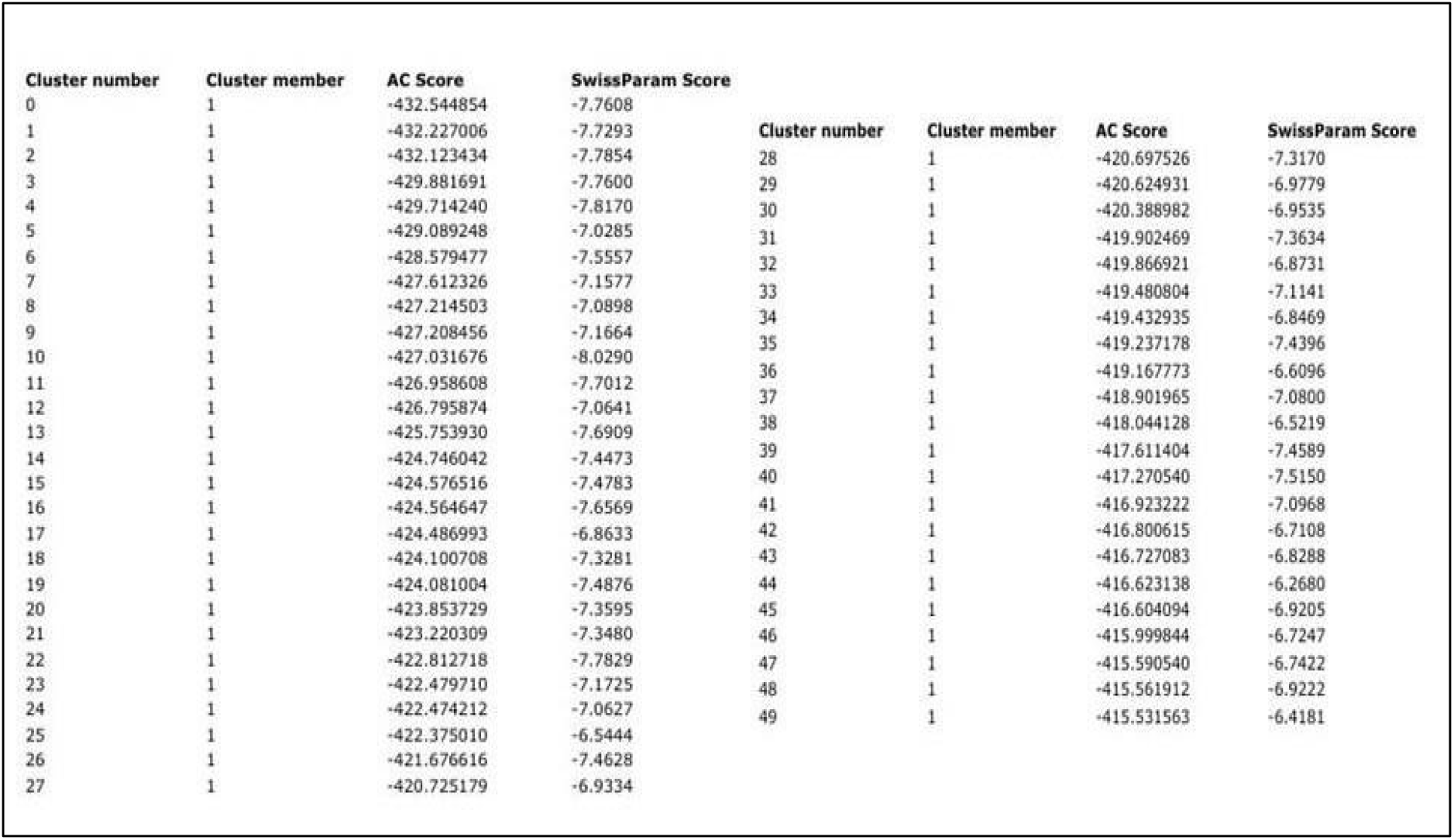
SwissDock docking scores for ATP binding to Pro1069, Ala1070 and Tyr1072 residues of AsCas12a.

**Table 5.** Possible ATP nucleotide binding site of AsCas12a according to ATPbind.

| Binding Pocket ID | Ligand | Binding Sites |
| --- | --- | --- |
| 1 (PAM-interacting domain) | ATP | Arg696, Gln700 |
| 2 (NUC domain) | ATP | Ser1068, His1071 |

**Figure 3.**
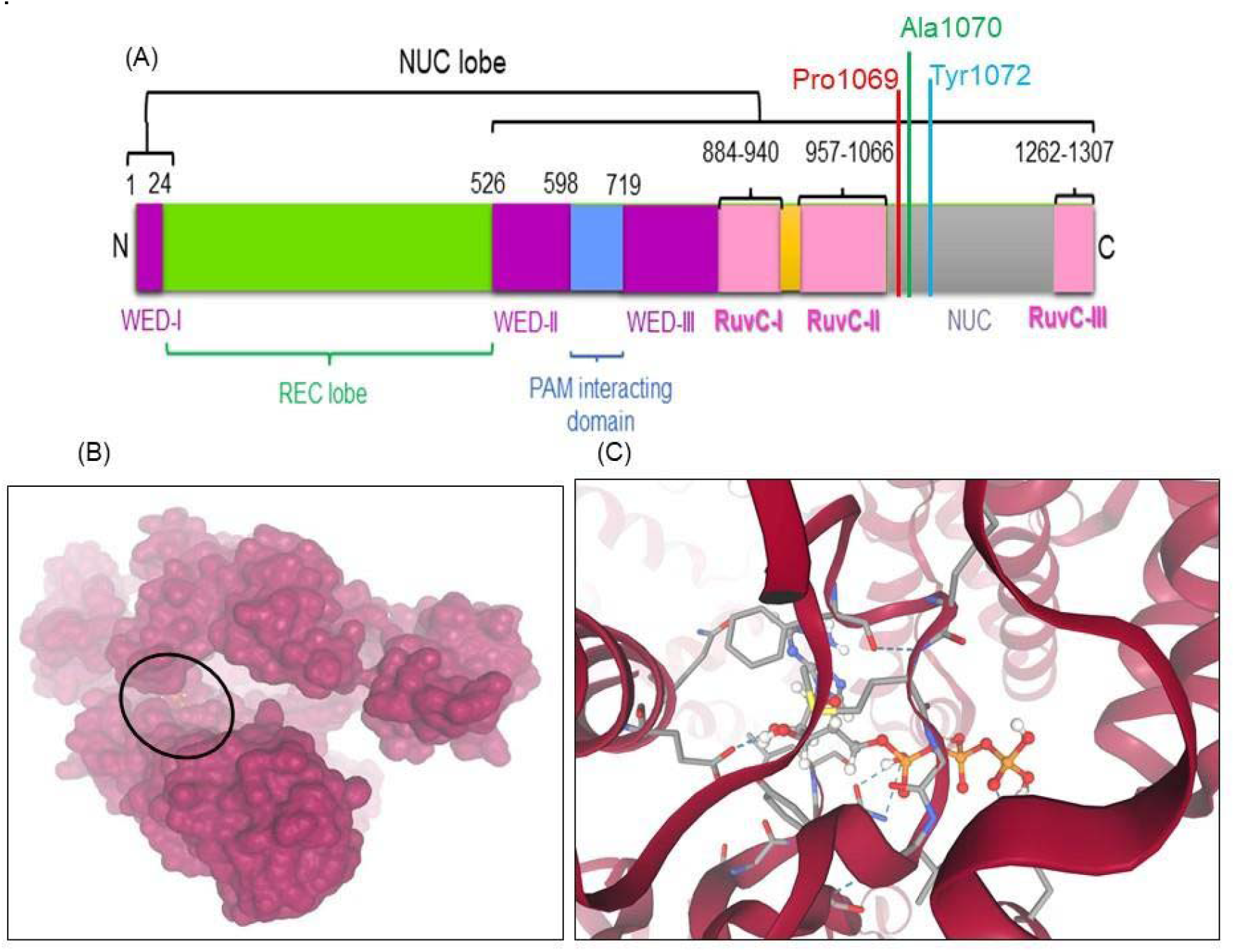
*In silico* prediction of ATP-binding residues in AsCas12a.(A) Schematic representation of the domain architecture of AsCas12a, with predicted ATP-binding residues indicated as distinctly colored bars within the NUC (nuclease) domain.(B) Surface rendering of the AsCas12a structure generated using SwissDock, illustrating the docked ATP molecule positioned deep within a defined region of the protein, marked by an oval. (C) Enlarged schematic view of the predicted ATP-binding pocket in AsCas12a, highlighting the spatial arrangement of the ATP molecule within the selected region.

**Figure 5.**
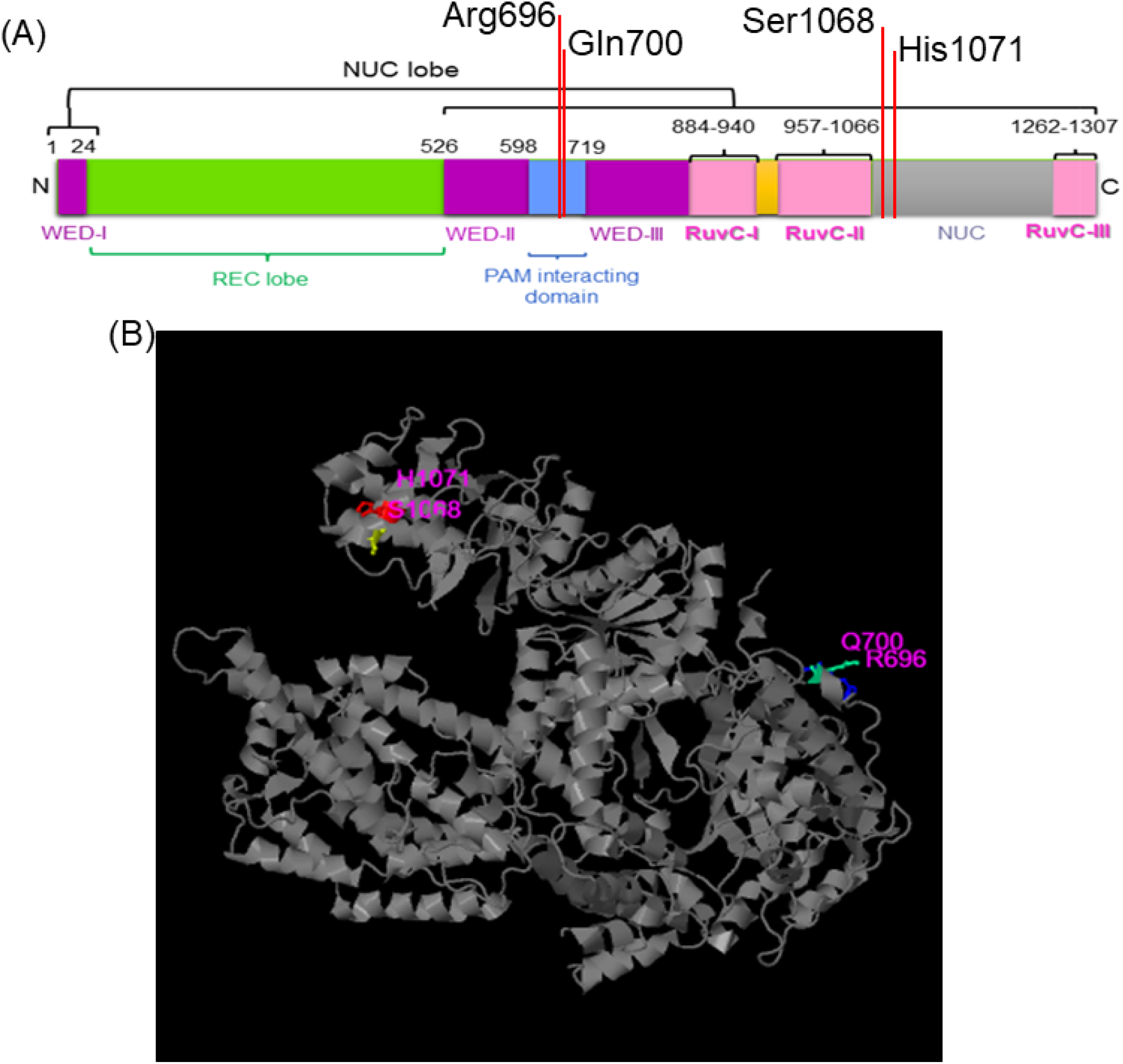
ATPbind predicts ATP-binding residues in AsCas12a. (A) Schematic representation of the domain architecture of AsCas12a, with predicted ATP-binding residues indicated as red-colored bars within the PI (PAM-interacting) and NUC (nuclease) domains. (B) AsCas12a structure generated using ATPbind, illustrating the ATP molecule positioned within the defined regions of the protein, and ATP interacting residues in magenta in standard one-letter amino acid code.

## 4. Discussion

Despite the absence of identifiable ATP-binding motifs, AsCas12a hydrolyzes ATP efficiently, and independently of DNA cofactors. This distinguishes it from classical DNA-dependent ATPases and raises intriguing mechanistic and functional questions. The biological relevance of this activity remains to be determined. It is possible that ATP hydrolysis could modulate conformational changes in Cas12a during its activation cycle or interactions with other cellular factors (17). Alternatively, the ATPase activity may reflect a vestigial or context-specific enzymatic function with regulatory implications.

Notably, ATPase activity is not a known requirement for Cas12a’s DNA cleavage function, which relies on a metal-dependent RuvC domain (18). However, auxiliary ATPase and ATP deamination activities in other CRISPR systems have been documented, particularly in Cas3 and CRISPR-associated Cad1 complexes, respectively, supporting the idea that CRISPR proteins may carry multifunctional capabilities (19, 20). Further biochemical, structural and *in vivo* studies will be necessary to elucidate the mechanistic basis and potential physiological role of this novel ATPase activity.

## Authorship contribution

SB: Investigation, Visualization, Methodology; Writing-final version.

## Declaration of competing interest

The author declares no competing interests are involved.

## Acknowledgements

I thank Osamu Nureki (University of Tokyo) for sharing the pE-AsCas12a expression plasmid. This work was funded the Indian Institute of Science fellowship to SB. I also thank the Department of Biochemistry, Indian Institute of Science for providing facilities to conduct protein purifications and biochemical assays.

## Abbreviations

ATP: Adenosine triphosphate;
CRISPR-Cas: Clustered regularly interspaced short palindromic repeats-CRISPR associated;
dsDNA: Double-stranded DNA;
IPTG: Isopropyl-β-D-thiogalactopyranoside;
ODN: oligonucleotide;
ssDNA: Single-stranded DNA

## Disclaimer

This study presents preliminary findings that have not yet been peer-reviewed. Feedback from the scientific community is welcome.

## Supplementary Appendix-

**Figure S1.**
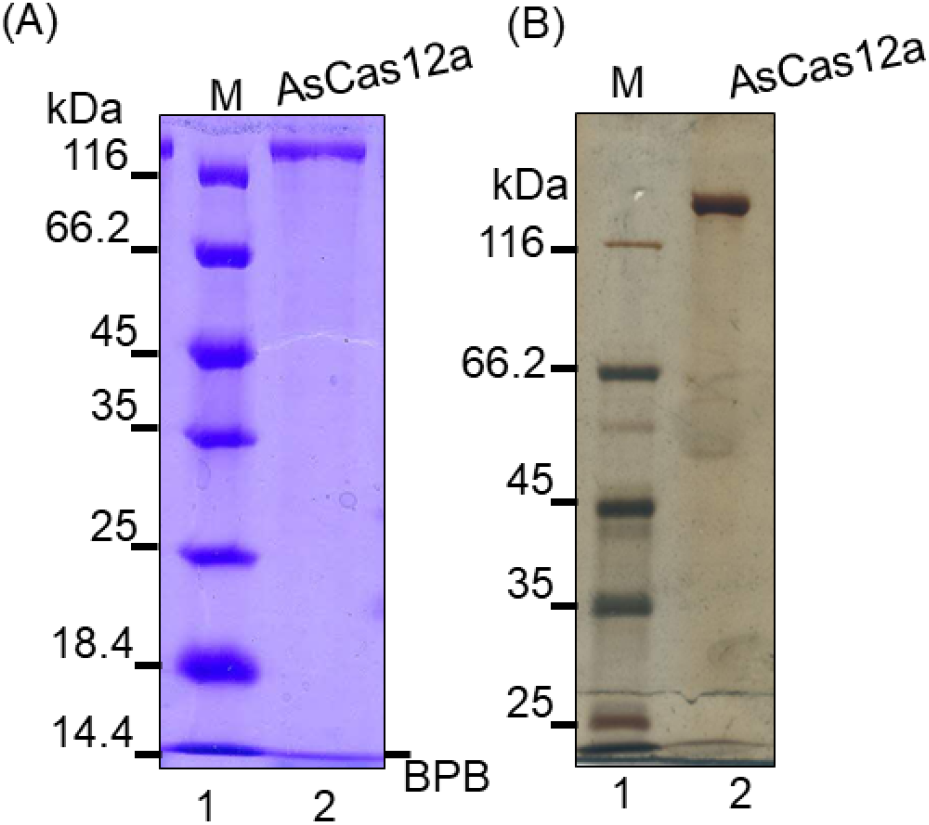
Purification profile of AsCas12a. 3 µg of AsCas12a was loaded onto a 7.5% SDS-PAGE gel, and stained with either Coomassie Brilliant Blue R-250 (A), or the gel was silver-stained (B) to assess for purity. Lane 1, molecular weight markers (M), Lane 2, AsCas12a alone. kDa-kilo-daltons, BPB-bromophenol blue.

## Notes

### Competing Interest Statement

The authors have declared no competing interest.

### Summary of Updates

This revision contains a prediction of ATP-binding sites using a new tool, ATPbind.

